# Neural and behavioral correlates of evidence accumulation in human click-based echolocation

**DOI:** 10.1101/2025.08.30.673202

**Authors:** Haydée G García-Lázaro, Santani Teng

**Affiliations:** Smith-Kettlewell Eye Research Institute

**Keywords:** echolocation, spatial localization, EEG, MVPA, psychophysics, blindness

## Abstract

Echolocation enables blind individuals to perceive and navigate their environment by emitting clicks and interpreting their returning echoes. While expert blind echolocators demonstrate remarkable spatial accuracy, the behavioral and neural mechanisms supporting the temporal integration of spatial echoacoustic cues remain less explored. Here, we investigated the temporal dynamics of spatial information accumulation in human click-based echolocation using EEG. Blind expert echolocators and novice sighted participants localized virtual spatialized echoes derived from realistic synthesized mouth clicks, with trials presenting trains of 2–11 clicks.

Behavioral results showed that blind expert echolocators significantly outperformed sighted controls in spatial localization. For these experts, localization thresholds decreased with more clicks, indicating cumulative integration of spatial cues across repeated samples. EEG decoding analyses revealed that neural representations significantly distinguished echo laterality and predicted overall spatial localization performance from the first click alone. Additionally, brain responses relative to the first click evolved systematically over successive clicks, paralleling psychophysical performance in blind echolocators and providing a possible index of perceptual information accumulation. These findings provide, to our knowledge, the first fine-grained account of temporal neural dynamics underlying click-based echolocation, directly linked to behavioral performance over multiple samples. They reveal how successive echoes are integrated over time into coherent spatial representations. Together, these results advance our understanding of the perceptual and neural mechanisms underlying echolocation and demonstrate adaptive sensory processing in the absence of vision.

## INTRODUCTION

Human echolocation is a remarkable form of active sensing that enables some blind individuals to perceive and navigate their environment using sound. By producing brief sounds, typically mouth clicks, and interpreting the echoes reflected from surrounding surfaces, echolocators extract spatial information to map their surroundings. Proficient blind echolocators consistently outperform non-expert blind and sighted individuals in various echoacoustic tasks. These include the perception of object distance, size, shape, texture, location, density, and motion (Hausfeld et al., 1982; Kellogg, 1962; Kolarik et al., 2014; Milne et al., 2014; Rice, 1969; B. Schenkman & Nilsson, 2010; B. N. Schenkman & Nilsson, 2011; Teng et al., 2011, 2024; Teng & Whitney, 2011; Thaler et al., 2013). The spatial resolution achieved by blind echolocators often surpasses that of sighted individuals (Teng & Whitney, 2011), even when their learning rates are comparable (Heller et al., 2017; Norman et al., 2021; Teng & Whitney, 2011; Tonelli et al., 2016). This distinction suggests that learning rate and spatial resolution likely reflect separate components of echolocation proficiency, shaped by different underlying mechanisms modulated by visual experience.

Beyond perceptual accuracy, proficient human echolocators dynamically modulate their sampling strategies to optimize spatial perception based on task demands and environmental complexity. For instance, they emit more and louder clicks under acoustically challenging conditions or when detecting and localizing objects at wider azimuths from the body midline (Norman & Thaler, 2018; Thaler et al., 2018, 2019), likely to enhance echo salience and signal-to-noise ratio. Target size also influences sampling behavior. When localizing smaller targets, echolocators engage in head-scanning movements that cover wider spatial areas, requiring more clicks and time before anchoring to near-target locations; in contrast, larger objects are typically localized with fewer clicks, reduced exploratory motion, and higher precision (Patel et al., 2024; Teng & Fusco, 2019). These adaptive behaviors suggest a flexible, goal-directed sampling process in which echolocators appear to gather more echoacoustic information over multiple clicks to refine their internal representations.

This dynamic modulation raises a fundamental computational question. Does spatial perception in echolocation rely on a single optimal echo, or is it the result of cumulative echoacoustic information gathered across multiple echoes? In audiovisual perception, sensory evidence accumulation frameworks describe how decisions emerge through the progressive integration of noisy inputs (Bondy et al., 2024; Hanks et al., 2015; Noppeney et al., 2010; Pinto et al., 2022; Yang et al., 2016; Yao et al., 2020). Whether a similar process governs echoacoustic spatial perception remains unknown. One possibility is that repeated clicks increase the chance of producing an optimal sample, which could occur at any point in a click sequence, without further benefit from repetitions and information integration. Alternatively, if cumulative echoacoustic integration is a fundamental process for echoacoustic perception, each additional sample should contribute incrementally to the refinement of a spatial percept. Distinguishing between these accounts is essential to understanding the principles by which humans extract spatial information through echolocation.

To address this question, here we investigated the neural and behavioral dynamics of echoacoustic spatial perception. Using electroencephalographic (EEG) recordings, we examined neural and behavioral responses while blind expert echolocators and sighted novice controls performed a spatial localization task requiring them to localize virtual objects based on spatialized echoes. Participants heard synthetic clicks and their spatialized reflections, simulating a virtual object located at varying eccentricities in the azimuth, and indicated whether the object was located to the left or right of the midline. We tested three primary hypotheses: (i) localization performance would improve systematically with increasing azimuthal eccentricity of the echo; (ii) localization performance would improve systematically with increasing numbers of echo-acoustic samples, consistent with evidence accumulation rather than optimal single-sample processing; and (iii) expert blind echolocators would demonstrate superior localization accuracy compared to sighted controls, with both behavioral and neural measures reflecting expertise-driven differences in spatial processing. Our findings confirmed both hypotheses. Localization accuracy improved with additional clicks in blind practitioners, expert blind echolocators outperformed sighted controls, and EEG decoding accuracy and variability across clicks reflect perceptual performance. Together, these results support the idea that spatial representations in echolocation evolve through the integration of cumulative sensory information over time, gated by the perceptibility of the individual echoes. These findings advance our understanding of human echolocation mechanisms and provide a basis for understanding cross-modal plasticity in sensory loss.

## METHODS

### Participants

In this study, twenty-one sighted individuals (mean age = 32.3 years, SD = 5.7 years, 12 males) and four blind expert echolocators (3 EB = early-blind and 1 LB = late-blind, all male, ages [16-58 y]) participated. Proficient echolocators were recruited based on self-reports of long-term and active use of mouth-clicking echolocation for daily life activities (see details in Table 1). Sighted controls were untrained in echolocation and reported normal or corrected-to-normal vision. Both sighted and blind individuals were naive to the task; they had normal hearing as assessed with pure tone audiometry (250-8000 Hz) and gave informed consent following our protocol approved by the Smith-Kettlewell Eye Research Institute Institutional Review Board.

**Table 1.** Blind expert echolocator participants.

| Blind individuals |  |  |  |  |  |
| --- | --- | --- | --- | --- | --- |
| Subject ID | Blindness onset | Age at test (y) | Blindness severity | Blindness Etiology | Echolocation expertise (# years of practice) |
| EB1 | 0y | 46 | Total | Glaucoma and optic nerve damage | >40 |
| EB2 | <1y | 57 | Total | Retinoblastoma (Enucleation) | ~57 |
| EB3 | 0y | 16 | LP | Retinal detachment | 11 |
| LB1 | 14y | 39 | Total | Optic nerve atrophy | 24 |

*LP= light perception*

### Stimuli

Stimuli consisted of a ∼6-ms convolved sound of a mouth click and its spatialized echoes reflecting a virtual object located 1 m away at various eccentricities in the azimuthal plane. To generate the stimuli, we first synthesized a mouth click using parameters and code from Thaler et al. (2017), modeling spectrotemporal waveform features and azimuthal directivity of clicks produced by skilled human echolocators (∼3 ms duration, power distributed from ∼1-15 kHz, peaking at ∼2-4 kHz). To simulate the spatial origin of the oral click, the waveform was spatialized at zero degrees (directly in the frontal plane) by convolution with a generic head-related transfer function (HRTF) (Gardner & Martin, 1995). The echo was simulated by introducing a time delay corresponding to the 1 m distance (∼5.8 ms at 343 m/s) plus an estimated 12 cm between mouth and ear, attenuation from geometric spreading loss, and the emitted pulse’s azimuthal directivity and spatialization via HRTF convolution at 5°, 10°, 15°, 20°, or 25° to the left or right of center. No losses from target reflectivity were modeled. The final binaural stimulus, comprising mouth clicks and echoes, was ∼6 ms in duration.

### Task and Experimental Procedure

The experiment was conducted in a darkened, sound-damped testing booth (Vocal Booth, Bend, OR). Participants sat 57 cm from a 27” display (Asus ROG Swift PG278QR, Asus, Taipei, Taiwan), wearing the EEG cap with 64 channels and tubal-insert earphones (Etymotic ER-3A). Stimuli were binaural and presented dichotically. We used the MOTU UltraLite Mk4 Audio Interface (MOTU Inc., Cambridge, MA) to improve the temporal precision of auditory stimulus presentation and synchronization to EEG triggers via pass-through detection (StimTrak, Brain Products Inc.). Auditory stimuli were presented at 64 dB SPL.

Participants completed a 2AFC auditory localization task in which they reported whether a virtual object was located to the left or right relative to the midsagittal plane. For sighted participants, each trial began with a white fixation cross against a gray display, followed by a sequence of 2, 5, 8, or 11 identical click-echo stimuli, separated by a stimulus-onset asynchrony (SOA) of 750 ms (Figure 1.B.3). The number of clicks was not indicated ahead of time. Following the final stimulus in the train, a 5-ms 180 Hz tone cued an untimed response display; subjects responded in two steps: cycling through “left” and “right” response options using the up and down arrow keys, then confirming with either of the adjacent (left/right arrow) keys. The initially displayed response option was randomized. This response scheme was made clear to subjects from the start of the experiment during the practice blocks. By jittering the sequence of keypresses needed for a given response in each trial, we decoupled stimulus-related processing from motor response planning and execution, as in previous work (García-Lázaro & Teng, 2024). A 5-ms 200-Hz tone confirmed the response and initiated a jittered intertrial interval (ITI) of 700–1250 ms before the next trial (Figure 1.B.3).

**Figure 1.**
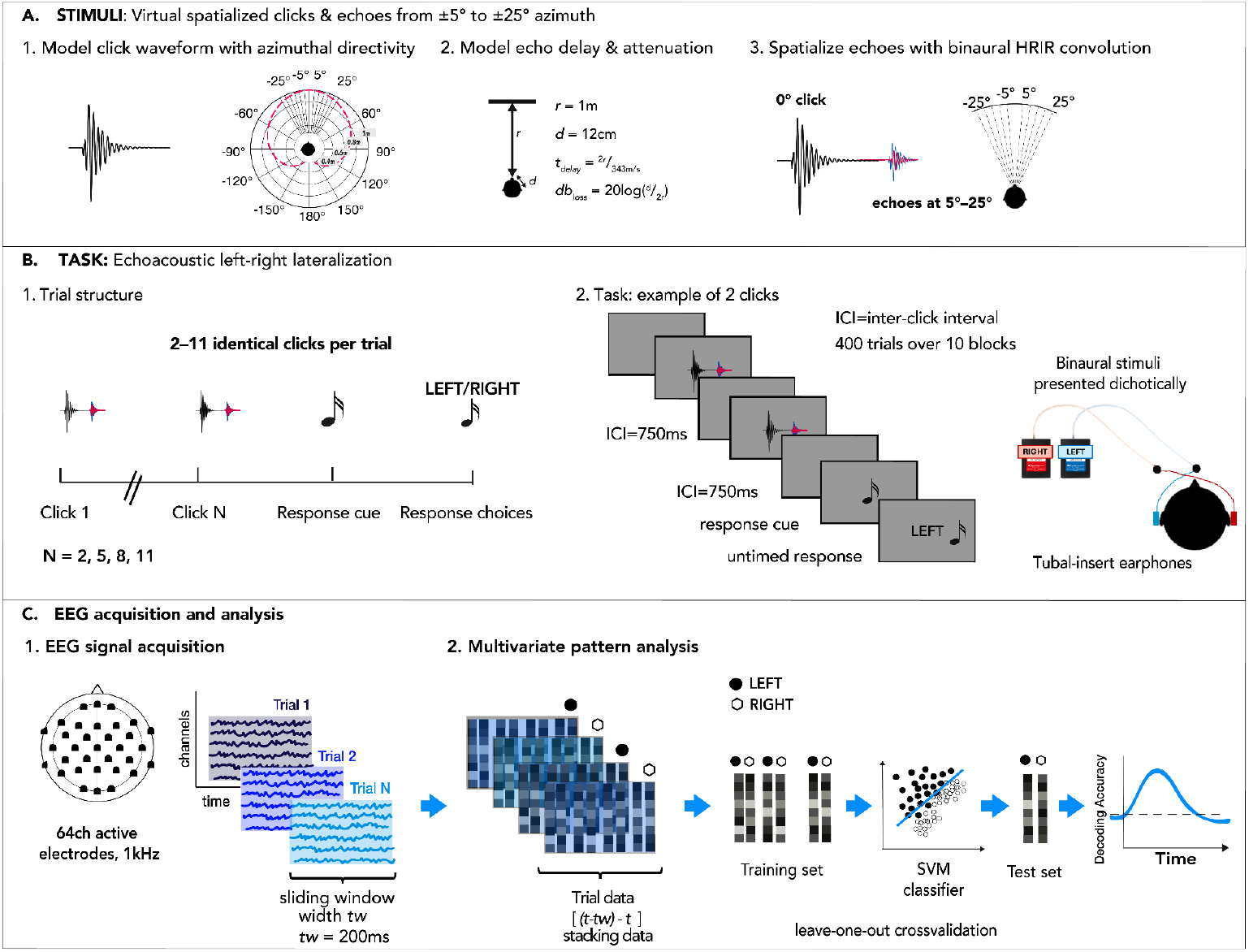
Stimulus generation, experimental design, and EEG acquisition/analysis. A. STIMULI: 1. Click generation and modeling azimuthal directivity (Thaler et al., 2017). 2. Echo delay and attenuation modeled based on 1-m distance. No reflectivity loss was modeled. 3. Echo spatialization (5°–25°) using head-related impulse response (HRIR) (Gardner & Martin, 1995). B. TASK: 1. Trials structured in trains of 2, 5, 8, or 11 clicks. 2. Task: an example of a trial of two clicks. C. EEG acquisition and analysis: 1. EEG signal acquisition. 2. Multivariate pattern analysis pipeline to generate decoding time courses.

We blocked trials into 10 groups of 40 trials each, presented in random order and corresponding to 40 conditions created by combining 10 eccentricities (±5°, ±10°, ±15°, ±20°, and ±25°) and 4 sequences of varying clicks (2, 5, 8, and 11). We collapsed left and right positions to focus on absolute eccentricity relative to the midline. Each block lasted ∼7 min, with breaks between blocks as needed, for a total experiment time of ∼70-75 min. We used Psychtoolbox-3 (Pelli, 1997) running in Matlab 2021b (The Mathworks, Natick, MA) for stimulus presentation, the Matlab Audio Processing Toolbox, and the voice changer tool (*Time-Domain Pitch and Time Scale Modification of Speech Signal*) to speed spoken instructions for blind individuals (1.8x normal speed) (Nguyen Tien Dung and Nguyen Dinh Hai, 2008).

### EEG data acquisition and preprocessing

We recorded continuous EEG using a Brain Products actiCHamp Plus recording system (Brain Products GmbH, Gilching, Germany) with 64 channels arranged in a modified 10-20 configuration on the caps (Easycap GmbH, Herrsching, Germany). The *Fz* channel was used as an online reference during the recording. We used the StimTrak device (Brain Products GmbH) as an auxiliary EEG channel connected to amplifiers and audio output earphones to tag stimulus onset times more precisely. The EEG signal was band-pass filtered online from 0.01 to 500 Hz and digitized at 1000 Hz. The continuous EEG signal was preprocessed offline using the FieldTrip toolbox (Oostenveld et al., 2011) and customized scripts using Matlab functions for downsampling and filtering the neural signal. Raw data were re-referenced to the common average of all electrodes and segmented into epochs of different lengths according to conditions. Stimulus onset times were corrected and aligned with the timestamps provided by StimTrak to increase time precision. Trials of the first 2 and last 2 clicks were segmented from -400 ms to 1400 ms relative to the stimulus onset of interest. Trials of 5 clicks were segmented from -400 to 3700 ms; trials of 8 clicks were from -400 to 5950 ms; and trials of 11 clicks were from -400 to 8200 ms. Epochs were baseline corrected, downsampled by averaging across non-overlapping 10-ms windows (Guggenmos et al., 2018), and low-pass filtered at 30 Hz. Trials were labeled by target laterality (*Right, Left*) and number of click-echoes (*2, 5, 8,* or *11 Clicks*).

#### EEG Multivariate Pattern Analysis (MVPA)

We analyzed the EEG signal using linear support vector machine (SVM) classifiers to decode neural response patterns at each time point of the preprocessed epoch using a multivariate pattern analysis (MVPA) approach (Figure 1.C). We applied a retrospective sliding window in which the classifier for time point t was trained with preprocessed and subsampled sensor-wise data in the interval [t-20, t]. This method increases the signal-to-noise ratio (SNR) and captures the temporal integration of dynamic and non-stationary properties (García-Lázaro &
 Teng, 2024). The resultant decoding time course thus began at -200 ms relative to stimulus onset. Pattern vectors in the window were stacked to make a composite vector. For example, 21 samples of 64-channel data made a composite vector that was 1344 elements long. Decoding was conducted using custom Matlab scripts that adapted functions from Brainstorm’s MVPA package (Tadel et al., 2011) and *libsvm* (Chang & Lin, 2011). We used 10-fold leave-one-out cross-validation, in which trials from each class were randomly assigned to 10 subsets and subaveraged (Guggenmos et al., 2018). This procedure was repeated with 100 permutations of subaverage sets; the final decoding accuracy for *t* represents the average across permutations.

### Statistical testing

To assess the statistical significance of behavioral indices, we used a one-sample binomial test to contrast the proportion of correct responses against chance (50%) for single-blind individuals and a t-test for the group of sighted controls. To compare the performance of blind echolocators with sighted controls, we used an adjusted one-tailed t-test analysis for small sample sizes as described by Crawford (2010) and Crawford and Howell (1998). The statistical significance of the EEG decoding time courses across sighted subjects was determined using t-tests against the null hypothesis of chance level (50%). We used non-parametric permutation-based cluster-size inference to control for error rate inflation in multiple comparisons. The cluster threshold was set to α = 0.05 (right-tailed) with 1000 permutations to create an empirical null hypothesis distribution. The permutation distribution’s significance probabilities and critical values were estimated using a Monte Carlo simulation (Maris & Oostenveld, 2007). The statistical significance of the EEG decoding time courses for single subjects was determined by contrasting them against a null distribution generated by randomly shuffling labels 100 times under the null hypothesis that left versus right labels were interchangeable. Decoding accuracy of the observed data was significant if it exceeded 95% of the permuted decoding scores.

## RESULTS

### Expert blind echolocators outperformed sighted controls in echolateralization

Overall performance for each echolocator and the group of sighted controls is shown in Fig. 2A. Blind echolocators were not all equally accurate, but all of them were better than chance at lateralizing the echoes (one-sample binomial test: EB1 mean = 99.5, p < 0.001; EB2 mean = 92.81, p < 0.001; EB3 mean = 88.25, p < 0.001; and LB1 mean = 56.5, p < 0.05; chance = 50). In contrast, sighted controls, as a group, performed at chance (SC mean = 51.83, s.e.m. = 0.98, t(20) = 1.86, *p* = 0.076). EBs’ performance compared to SC’s performance using an adjusted one-tailed t-test analysis for small samples (Crawford et al., 2010; Crawford & Howell, 1998) revealed that EBs, on average, differed from sighted controls (EBs’ mean = 93.52; SC mean = 51.83, SD = 4.50; t = 8.04, *p* < 0.001). LB performance did not statistically differ from SC performance (t = 1.04, *p* = 0.16).

**Figure 2.**
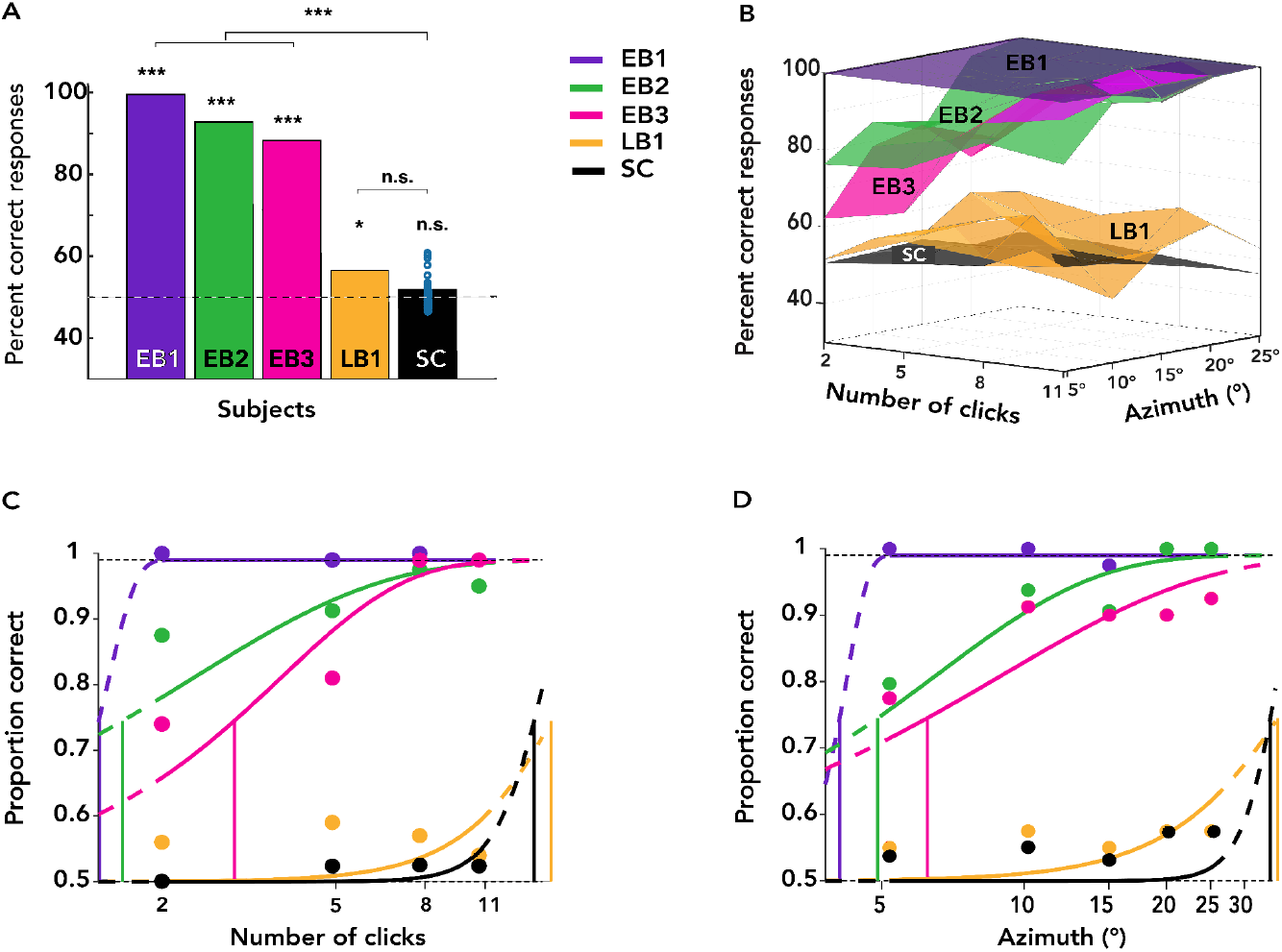
Task performance from early blind (EB), late blind (LB), and sighted control (SC) participants. A: Individual task performance according to visual experience. B: 3D visualization of task performance for azimuth and number of clicks. C-D. Psychometric functions for number of clicks and azimuth for each blind echolocator and the average of sighted controls (N = 21).

Figure 2B plots a 3D visualization of condition-wise performance for each individual and group, with the x-axis indicating click counts from 2 to 11 and the y-axis indicating absolute azimuthal eccentricities from 5° to 25°. We observed individual differences in echolocation performance between experts; for example, EB1 (violet) performed near ceiling in the task, while LB1 did not exceed 75% in any condition. In contrast, sighted controls did not perform higher than 61% when split by the number of clicks or eccentricities.

### Performance improved with azimuth and clicks, varying by expertise

To further investigate the role of click count and azimuthal eccentricity in echolateralization, we computed psychophysical thresholds for each blind participant and pooled sighted controls across all trials, grouping scores by the number of clicks or azimuth angles. The underlying psychometric curve was modeled using the Weibull function (Teng et al., 2012), with thresholds defined as the stimulus eccentricity or click count at which participants correctly lateralized echoes 75% of the time. The lapse rate was fixed at 0.01, with slope, width, and threshold set as free parameters. We used the psignifit4 toolbox (Schütt et al., 2016) in MATLAB 2021b for these analyses.

Figures 2C-D display separate psychometric functions for each EB and LB participant, along with a combined psychometric fit for SC, for azimuth and number of clicks. For EB participants, the proportion of correct responses improved with both increased azimuth degrees and the number of clicks. Thresholds for azimuth and number of clicks, respectively, varied across participants: EB1 (3.89°; 1.42 clicks; violet), EB2 (4.71°; 1.62; green), and EB3 (6.03°; 2.95; pink). In contrast, LB’s and SC’s chance or near-chance performance resulted in thresholds beyond the range of the sampled stimulus space for both azimuth and number of clicks: LB1 (34.88, 16.21; yellow) and SC (33.57, 14.80; black). As a result, subsequent analyses focus on the three EB participants.

### Azimuthal thresholds decreased with additional clicks in EB echolocators

Next, to assess the effect of repeated clicks on localization performance, we computed separate azimuthal thresholds for trials grouped by click count. Figure 3A-C shows individual psychometric fits for EB1, EB2, and EB3 for trials with 2, 5, 8, and 11 clicks, indicated by gradually lighter shades of blue. Individual thresholds by click count are plotted in Figure 3D. For EB2 and EB3, azimuthal thresholds decreased with increasing number of clicks, reaching a plateau after 8 clicks for EB2 and EB3. In contrast, EB1’s ceiling-level thresholds were consistent regardless of the number of clicks. This indicates that EB2 and EB3 benefited from repeated identical click presentations, while EB1 needed no more than two clicks to perfectly lateralize echoes within a range of 5° to 25°. EB1’s ceiling-level thresholds also serve as a benchmark for performance limits in this experiment, indicating that EB2 and EB3 plateaued after 8 clicks.

**Figure 3.**
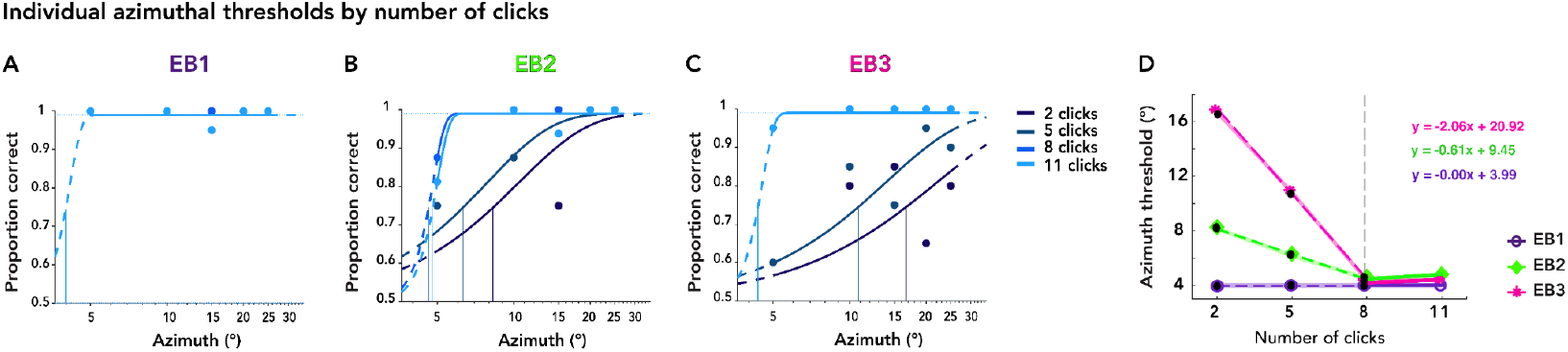
Individual psychometric functions for azimuth across click-echo conditions in early blind echolocators. A-C: Psychometric curves are shown for each early blind participant, illustrating localization performance as a function of azimuth angle, separately for different numbers of clicks. Vertical lines indicate 75% thresholds. D: Thresholds plotted against the number of click-echoes. Piecewise linear regression (dashed lines) is used to assess the change in spatial precision across the first three click-count conditions before the ceiling-performance plateau.

To quantify performance improvements with repeated clicks, we fitted piecewise linear models to the first three data points for each participant to capture the initial change rate before the plateau. The degree of improvement varied across individuals, reflected both in the initial threshold with two clicks and the slope of improvement with additional clicks, as captured by the model’s parameters. Figure 3D illustrates the linear fit for EB2: the model predicted an azimuthal precision of approximately ∼8° with two clicks, with each additional click improving precision by about 0.61° (threshold = –0.61 × clicks + 9.45). For EB3, the model estimated a higher starting threshold of ∼21°, with a steeper improvement of ∼2° per additional click (threshold = –2.06 × clicks + 20.42). EB1 already exhibited low thresholds with two clicks (∼3.99°) with no additional gain (but also no loss) with further repetitions.

Together, these results highlight a progressive improvement in spatial precision as the number of clicks increases, quantifying the estimated rate of information gain for each individual.

### Effect of azimuth on click-based thresholds for object spatial location

We next examined how azimuthal eccentricities related to the number of clicks required for accurate spatial localization. We grouped trials by azimuth angle to fit individual psychometric functions for the number of clicks. Figure 4 displays these psychometric functions for EB1, EB2, and EB3 with azimuths ranging from 5° (dark red) to 25° (light red) in increments of 5°. The threshold patterns shown in Figures 4 A-C reveal distinct individual differences across azimuths. For objects located at 5° of eccentricity, EB2 and EB3 required approximately 5.5 to 6.5 clicks to localize accurately. Thresholds declined sharply from 5° to 10° and then more gradually from 10° to 25°, suggesting that beyond 10° of eccentricity, EB2 and EB3 needed 1.5–3 clicks to localize objects. Notably, EB1 thresholds quickly reached 1.5 and remained steady at it across all azimuths, indicating that 2 clicks provided sufficient information to localize objects from the entire 5° to 25° range. These findings reveal individual differences in the number of clicks required to achieve spatial precision and illustrate that laterality information varies with eccentricity: a less eccentric echo has “less” laterality information that may require more clicks to perceive.

**Figure 4.**
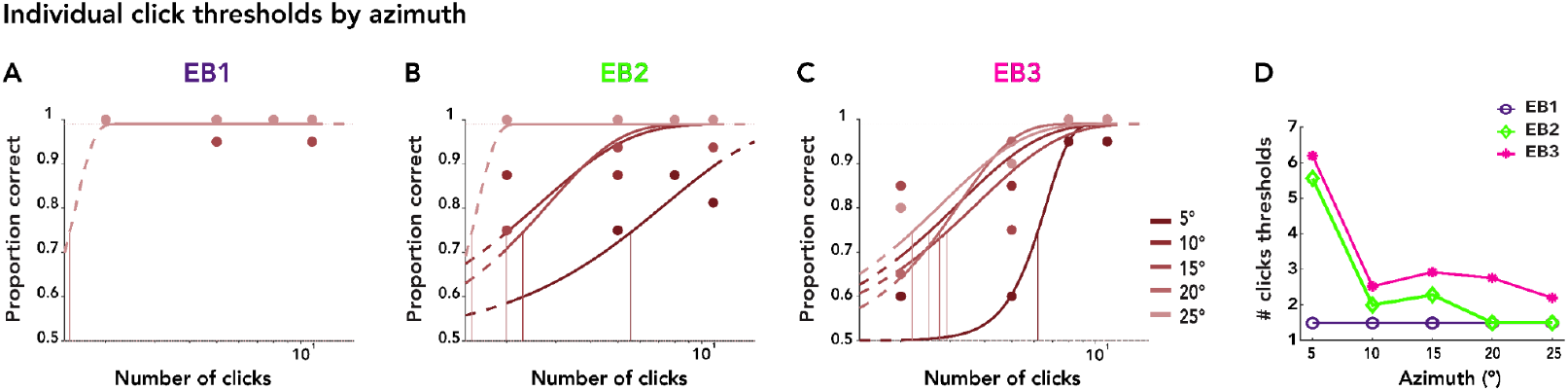
Individual psychometric functions for each early blind echolocator, for the number of clicks as the stimulus level, grouped by azimuth. A-C: Psychometric curves illustrate localization performance as a function of the number of clicks, plotted separately for each azimuth condition (red). D: Thresholds (derived from psychometric functions) are plotted against azimuth.

### Neural representations of spatial object laterality across early and late clicks

Next, we examined how echoacoustic spatial information is represented in neural activity by applying multivariate pattern analysis (MVPA) to EEG data. To capture the temporal evolution of spatial representations, we classified neural responses between left- and right-lateralized echoes, focusing on two time windows within each trial: the first two and the final two clicks. As each trial contained at least two clicks, this analysis allowed us to include all trials. More importantly, because the physical stimulation, participant sample, trial count, etc., were identical across these windows, differences in decoding dynamics were likely to arise from changes in the brain state across the click train rather than in the stimuli.

### Neural spatial encoding during early clicks

Figure 5A shows decoding time courses for individual participants and the pooled SC group, color-coded as in Fig. 2. In EB1 (violet), decoding accuracy rose sharply around 180 ms after the first click, peaked at 98%, dipped slightly around 600 ms, and then rose again following the second click, remaining elevated through the end of the epoch. Similarly, EB2 (green) exhibited above-chance decoding from 240–800 ms and 930–1400 ms, while EB3 (pink) demonstrated significant decoding from 70–990 ms. All EB participants exhibited robust and sustained decoding accuracy following both clicks, with peak decoding accuracy consistently occurring after the first click and a secondary peak following the second click. In contrast, LB1 (orange) showed above-chance decoding from 510 to 1060 ms, with peak accuracy following the second click. Sighted Controls (black) showed no significant decoding throughout the epoch. Notably, decoding peaks during the first two clicks perfectly match the rank order of behavioral performance.

**Figure 5.**
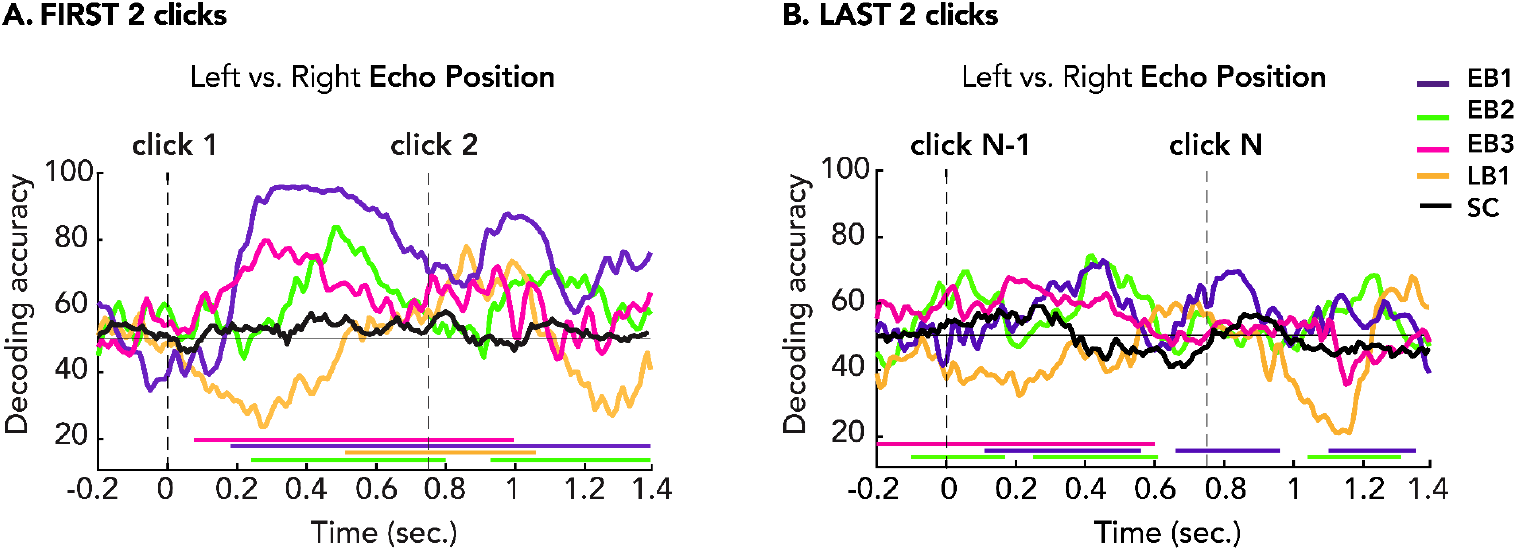
Decoding time courses for echo laterality following the first (A) and last two clicks (B) of each trial. Vertical dashed lines indicate click onsets at 0 and 750 ms. Horizontal dashed line indicates chance level (50%), and colored bars along the x-axis denote significance. For sighted controls (SC), group-level statistics were computed (N = 21) using a t-test against chance (50%), with a cluster-defining threshold of *p* < 0.05 and 1000 permutations. For blind individuals, single-subject statistics were computed by comparing each participant’s decoding curve to an empirical null distribution generated by applying SVM decoding to label-randomized trials, permuted 100 times.

### Neural response of spatial encoding during final clicks

Next, we analyzed neural activity during the last two clicks of each trial to examine how spatial representations evolved toward the end of the trial before preparing for motor responses. As shown in Figure 5B, EB1 (violet) exhibited significant decoding accuracy (DA) in three distinct time windows: 110–560 ms, 660–960 ms, and 1100–1360 ms. EB2 (green) showed above-chance decoding from -100 to 170 ms, 250–610 ms, and 1040–1310 ms. EB3 (pink) demonstrated significant decoding from -200 to 600 ms. In contrast, LB1 (orange) and sighted controls (black) did not exhibit above-chance decoding throughout the final two clicks. All decoding peaks in the late-click epoch were lower compared to those in the early-click epoch. EB2 and EB3 decoding curves were significantly positive in the baseline preceding the onset of the penultimate click. Notably, these were the same two participants whose behavioral performance improved for trials with more than 2 clicks (Fig. 3).

### Neural signals from blind expert echolocators reliably track spatial location representation throughout the trials

To further examine whether these discriminative neural patterns varied across different stages of the click sequence, we applied SVM classification to single-click segments between different ordinal positions in the click train. We reasoned that if the brain accumulates perceptually relevant echoacoustic information, the brain’s response to a click should change systematically across identical click repetitions. For example, representational changes separate from the sensory response should be decodable in the prestimulus baseline. Thus, we decoded the first click against the last click in each condition (1 vs. 2, 1 vs. 5, 1 vs. 8, and 1 vs. 11). Each 1-click epoch spanned -200 ms to 550 ms relative to click onset.

Between-ordinal-position click decoding results are shown for each blind echolocator in Figure 6A–D. A similar overall pattern emerged: Click 1 was significantly (and almost perfectly) distinguishable from Click 2 at every time point in each epoch, then progressively less so for Clicks 5, 8, and 11. The preclick baseline became progressively less discriminable (that is, more similar) for each click relative to the first (Fig. 6E). For example, EB1’s Click 1 baseline was perfectly distinguishable from Click 2, ∼70% decodable from Click 5, and not significantly distinguishable from Clicks 8 and 11. For EB2 and EB3, the Click 1 baseline was decodable compared to subsequent clicks until reaching nonsignificance for Click 11. For LB1, only the Click 2 baseline was decodable relative to Click 1. For all echolocators, all post-click brain responses for Clicks 2, 5, 8, and 11 were significantly decodable from Click 1 by 100 ms and remained so for the rest of the epoch.

**Figure 6.**
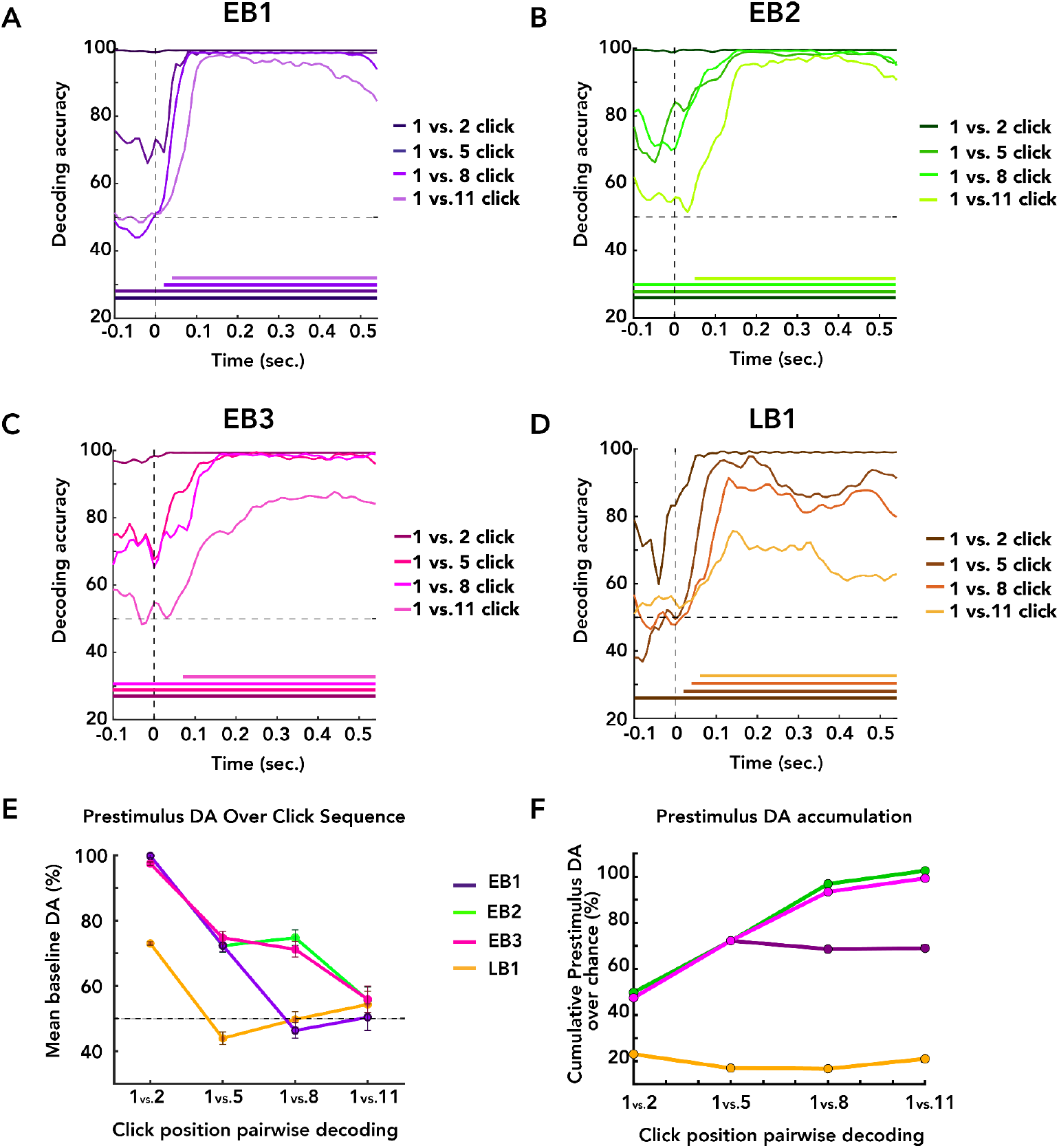
Neural decoding across click sequences in blind echolocators. A–D: Decoding accuracy time courses for individual blind echolocators. Colored lines indicate SVM classification contrasting the first click with the last click in each condition (1 vs. 2, 1 vs. 5, 1 vs. 8, 1 vs. 11). Each click was analyzed as a 1-click epoch from –200 to 550 ms relative to click onset. Horizontal bars along the x-axis mark time windows of significant above-chance decoding accuracy (DA) (null permutation; cluster-corrected, *p* < 0.05). E: Mean and standard deviation of DA from the baseline period between click conditions for each participant. DA was highest for initial click comparisons (1 vs. 2) and declined thereafter. F: Cumulative sum of DAs from Panel E, over chance level. Rising slope suggests increasing information over clicks; plateaus suggest plateau or failure to integrate.

## DISCUSSION

In this study, we investigated the neural and behavioral mechanisms underlying echoacoustic spatial perception by combining psychophysics with EEG recordings in blind expert echolocators and sighted novice controls. Participants performed a spatial lateralization task using virtual echoes derived from synthetic clicks, allowing us to examine how the human brain integrates temporally distributed auditory cues to construct spatial representations. Localization accuracy improved systematically with both eccentricity and the number of echoacoustic samples, consistent with a temporal evidence accumulation process. Expert blind echolocators exhibited significantly greater accuracy than novice sighted controls, suggesting experience-dependent differences in spatial processing. Neural decoding analyses closely mirrored behavioral performance, suggesting that dynamic neural representations reflected the evolving spatial perception. These findings demonstrate that spatial representations in echolocation are constructed through the progressive integration of discrete echoacoustic samples, contingent on echo perceptibility.

### Expert blind echolocators exhibit superior spatial localization, shaped by visual experience

Consistent with prior research findings (Rice, 1967, 1969; Vercillo et al., 2015, 2017; Wallmeier et al., 2015), blind expert echolocators tended to outperform sighted controls in localizing objects. Although all blind echolocators performed above chance, EB individuals were more accurate, performing at >70% even with only 2 clicks. LB’s lower performance comes despite being proficient in real-world echolocation and serving as an echolocation instructor. This performance difference based on blindness onset aligns with findings by Thaler et al. (2011), who reported lower performance in an LB expert compared to EB, including chance-level lateralization of passively presented click-echo recordings. More broadly, the behavioral results are consistent with the idea that blindness onset critically shapes spatial hearing. Early blindness may foster auditory-based spatial priors and enhanced sensitivity to binaural cues (Nilsson & Schenkman, 2016; Röder et al., 1999), while later blindness leaves individuals relying on visually calibrated reference frames less suited to auditory judgments such as echolocation (Amadeo et al., 2019). This aligns with broader evidence that spatial perception is visually calibrated (Eimer, 2004; Gori et al., 2014; Röder et al., 2008; Röder & Kekunnaya, 2021; Sourav et al., 2018) and suggests that early deprivation coupled with echolocation enhances sensitivity to spatial acoustic cues and supports more efficient neural mechanisms for spatial decisions (Vercillo et al., 2017).

### Psychophysical thresholds reveal temporal evidence accumulation and spatial cue salience effects

Localization performance improved with increasing azimuthal eccentricity and additional clicks, especially in early-blind participants. These findings align with an evidence-accumulation account in which spatial information is scaled by eccentricity and integrated over repeated samples to drive perceptual decisions. The effect of azimuth likely reflects greater cue discriminability and enhanced interaural differences, supporting the notion that cue strength and salience jointly modulate the rate of evidence accumulation. This is corroborated by the click count thresholds in Fig. 4, showing that fewer clicks were needed to reach threshold performance when the echoes were more eccentric. We note that click acoustic energy is nearly isotropic within the range of our azimuths (Thaler et al., 2017), but that diminishing emission energy may offset the benefits of interaural cues at greater eccentricities. Recent work suggests an acoustically optimal eccentricity for echolocalization of about 45° (Thaler et al., 2022).

From a signal detection perspective, multiple echoes increase the signal-to-noise ratio by providing convergent information that boosts perceptual certainty (L. Thaler et al. 2018). Behaviorally, this process was captured by progressively decreasing lateralization thresholds with successive clicks in blind echolocators (Fig. 2). This pattern was clearest in EB2 and EB3, who showed steep improvements between the first and eighth clicks, suggesting their perceptual system effectively integrates echoacoustic features over time, then plateaus or saturates as ceiling performance is reached. (EB1’s thresholds did not improve with more clicks, as they were at ceiling level for all conditions.) The linear progression of thresholds provides a quantitative estimate of each individual click’s contribution to precision: about 0.6° per click for EB2, and 2° per click for EB3 (Fig. 3D). It also tentatively predicts a single-click lateralization threshold — 8.8° for EB2 and 18.9° for EB3. Figs. 2B-C also show that in addition to the threshold changes, EB2’s and EB3’s psychometric functions narrowed with more clicks, consistent with reduced uncertainty underlying the perceptual decision (Wichmann & Hill, 2001). LB1 performed above chance, but neither he nor sighted novices improved reliably across azimuths and click repetitions, suggesting that they were unable to extract and integrate meaningful spatial information from echoacoustic cues. Both LB1 individually and SC generally have been shown to echolocate successfully in other contexts (Norman et al., 2021, 2024; Teng & Whitney, 2011; Thaler et al., 2011), so some of the performance differences we show may be partly attributable to our experimental setup. We also note that our task probed finer spatial grain (5–25°) than some previous studies by Rice (0–90°) (1967, 1969), Thaler et al. (0–180°) (2018), and Wallmeier et al. (30–60°) (2015).

Echoacoustic information can accumulate dynamically in multiple ways, e.g., through the greater SNR of louder clicks (Thaler et al., 2018, 2019), integrating across viewpoints for shape (Milne et al., 2014; Teng et al., 2024), or integrating across positions for moving objects (Salles et al., 2020; Thaler et al., 2013). Here, by synthesizing the echoacoustic stimuli, we kept acoustic stimulation and target-observer relationship constant across clicks and participants. Our psychophysical results thus suggest that variation in click- and azimuth-dependent improvement across individuals reflects differences in baseline abilities or processing strategies, rather than fluctuating sensory input or idiosyncratic strategies. The behavioral improvements with more clicks are also more consistent with an information-accumulation model of perceptual decision-making, where each echo contributes discrete evidence toward resolving spatial uncertainty, rather than a foraging-like search for a single optimal sensory signal.

### Neural signals encode spatial location early and dynamically in blind echolocators

While numerous studies have investigated the brain mechanisms of echolocation via fMRI (Flanagin et al., 2017; Norman et al., 2024; Norman & Thaler, 2019; Thaler et al., 2011; Wallmeier et al., 2015), to our knowledge, the present study is the first to examine click-to-click neural dynamics during echoacoustic perception. EEG decoding timecourses revealed robust early encoding of echo laterality in blind expert echolocators. These neural representations were evident from the first clicks, with all EBs and LB1 showing significant decoding in the EEG response before the second click onset, although LB1’s response peaked afterward (Fig. 5A). Thus, interestingly, echo position is decodable in the EEG response after a single click. Because our behavioral probes started at two clicks, the decoding significance onset does not tell us whether that information was perceptually available to the observers. However, decoding peaks in the first-click epoch showed perfect rank-order agreement with overall behavioral accuracy for all four blind echolocators and the SC group (Spearman ρ = 1.0; Fig. 7). While technically significant despite the small sample size, this correlation supports a more descriptive observation in context with our other results: the initial brain response to a single click may index some generally predictive echolocation ability, even though performance did improve with additional clicks for the EBs.

**Figure 7.**
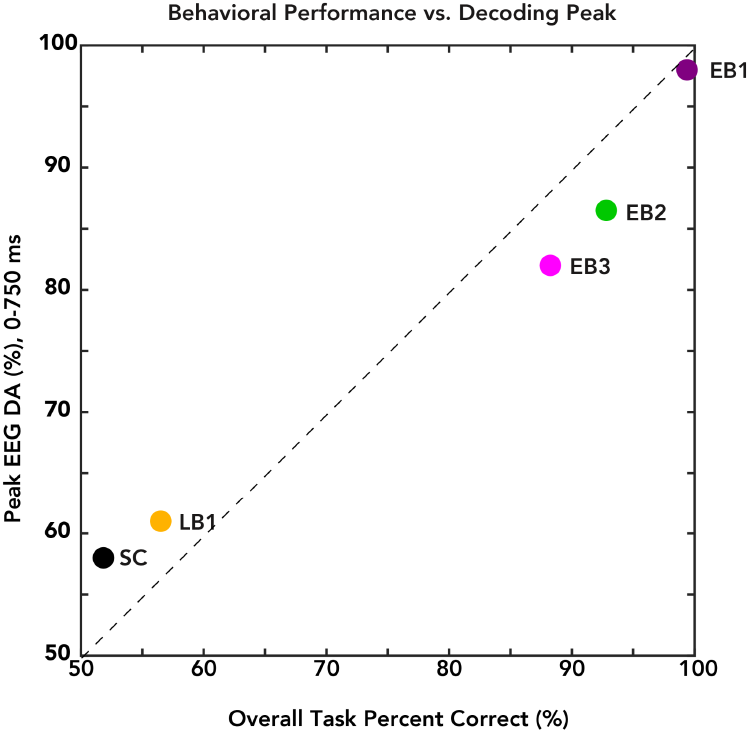
Correlation between the first DA peak in the 0-750 ms window (y-axis) and overall task performance (x-axis). Early blind participants (EB1–3) exhibit both higher decoding peaks and greater behavioral accuracy, whereas the late blind participant (LB1) and the sighted control (SC) show lower values. This pattern suggests that strong early spatial neural decoding predicts superior task performance. (Note: SC performance and decoding do not exceed chance level.)

Decoding of the last two clicks (Fig. 5B) showed persistent decodable responses for EBs, though all peaks were attenuated compared to the first two-clicks epoch. The inclusion of all trials in both analyses in Fig. 5 equates statistical power as well as the raw sensory stimulation—two acoustically identical clicks, equally spaced, in each epoch. The attenuated response is unsurprising despite the unchanging sensory information, attributable to well-known repetition suppression (RS) effects (Summerfield et al., 2011). The RS effect could mask any positive accumulation signals from being evident in the position decoding results. Still, the preclick positive decoding for EB2 and EB3 suggests that a persistent spatial representation is present near the tail of the click train that was absent at the beginning.

### Progressive evolution of the brain state over the click sequence

The decoding analysis between click ordinal positions in Fig. 6 accounts for RS to gauge how similar successive clicks are to the first click. The Click 1 vs. Click 2 decoding would theoretically index the largest state change in the brain about stimulus position, from zero to full acoustic information. In line with this prediction, we found Click 1 vs. 2 decoding accuracy to be the highest (e.g., nearly perfect across the entire click epoch for all EBs). Notably, over additional clicks, decoding dissimilarity to Click 1 evolved differently in the preclick compared to post-click periods. The preclick periods are the furthest removed from the sensory processing response and directly comparable with the prestimulus baseline. Preclick decoding across the click sequence therefore estimates how different the EEG response is from a prestimulus baseline when it is not actively processing sensory input. Applying this approach, we found that preclick DA compared to Click 1 decreased steadily after the initial maximum. For LB1 and EBs, the preclick period was indistinguishable from the Click 1 baseline by Click 5, 8, and 11, variously, even as the sensory response remained robustly decodable for every individual. Taking the above-chance preclick decodability from the prestimulus baseline as a measure of information available after each click, summing them yielded the pattern shown in Fig. 6F. Importantly, this pattern closely tracks the psychophysical threshold progressions and plateaus observed in the EBs (Fig. 3): EB1 “accumulates” before flattening after 5 clicks; EB2 and EB3 accumulate steadily until flattening after 8 clicks; and LB1 does not accumulate across clicks. In sum, the cross-click sequential decoding results may thus reveal how neural spatial representations evolve and accumulate over the click sequence.

### Potential of a sequence- and time-resolved approach to future work

The temporal dynamics observed in blind echolocators include a variety of signal trends: a strong initial response; attenuating decoding for each click, as well as signatures of persistent and accumulating representations across clicks. These may reflect dissociable signatures of sensory- and decision-related processing, a nested cascade in which echoacoustic features are rapidly extracted, integrated, and refined into coherent spatial percepts. Future research using multimodal neuroimaging could combine spatial and temporal precision (Cichy & Oliva, 2020; Lowe et al., 2022; Ritter & Villringer, 2006) to help identify the specific cortical circuits involved and clarify how these networks evolve through training and experience. Our time-resolved approach may also be applied to studies of more complex echo scenes and objects, better approximating real-world scenarios, and eventually integrate more ecologically valid aspects such as head movements or full-body mobility in virtualized environments (Gramann et al., 2010; Miyakoshi et al., 2021). Finally, granular examination of sequential click processing dynamics would help identify behavioral and neural underpinnings of expertise, moving past presumptions based on visual experience and self-report. As sighted observers can readily perform (or learn) echolocation to various extents, characterizing echolocation expertise in terms of sensitivity to spatial acoustic cues, efficiency of integration across time/samples, scene-analytic processing, etc. can more rigorously inform training interventions (García-Lázaro & Teng, 2025; Norman et al., 2021) and serves as a promising avenue to revealing generalizable principles of perceptual learning and sensory compensation.

### Summary and conclusion

The present study provides the first behavioral and neural characterization of individual click dynamics and their integration over repeated samples. The enhanced localization performance and early neural encoding in early blind echolocators reflect experience-dependent cortical adaptation to sensory deprivation. Our results may inform training programs aimed at enhancing echolocation in blind novices, as well as the development of assistive technologies that leverage temporal sampling strategies to support spatial perception. More fundamentally, this work highlights evidence accumulation as a key computational principle in human echolocation, showcases the remarkable flexibility of the brain’s perceptual systems in the absence of vision, and demonstrates a novel paradigm for future echolocation research.

## ACKNOWLEDGMENTS

This work was supported by Smith-Kettlewell Eye Research Institute, the Foundation for Ophthalmology Research and Education International (S.T.), the E. Matilda Ziegler Foundation for the Blind (S.T., H.G.L.) and Smith-Kettlewell Eye Research Institute’s C.V. Starr Fellowship Fund (H.G.L.). We thank our participants for their generosity with their time and feedback.

## AUTHOR CONTRIBUTIONS

Designed research: H.G.L.; S.T.; Performed research H.G.L., S.T.; Analyzed data: H.G.L., S.T.; Wrote the paper: H.G.L., S.T.

